## Supplementary Figs 1-5 and Table 1 for "Improved modeling of human vision by incorporating robustness to blur in convolutional neural networks"

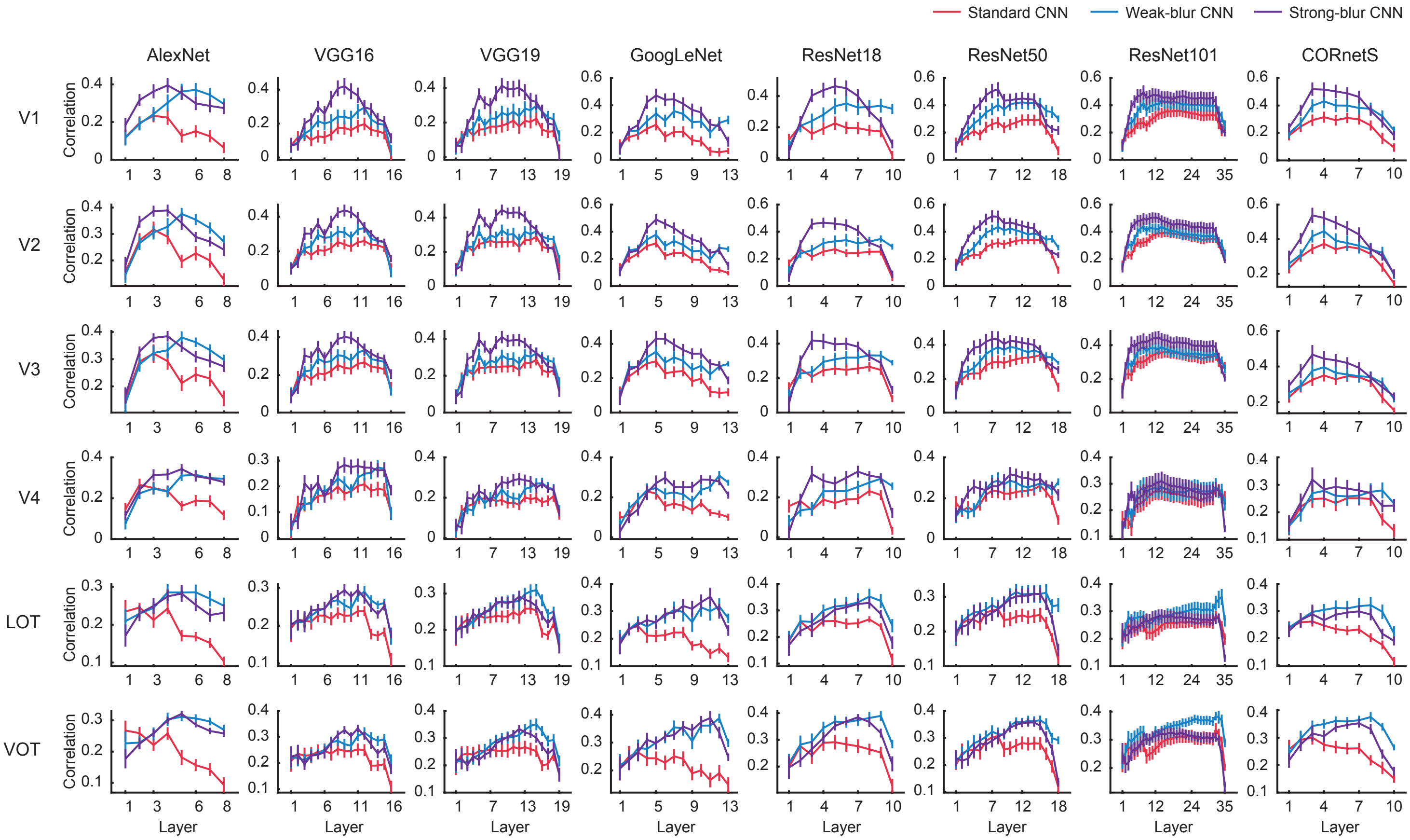


**Supplementary Figure 1.** Correlational similarity between CNN model responses and human neural responses to all viewing conditions, plotted for every CNN model by layer and for each visual area provided by Xu & Pashkam (2021).

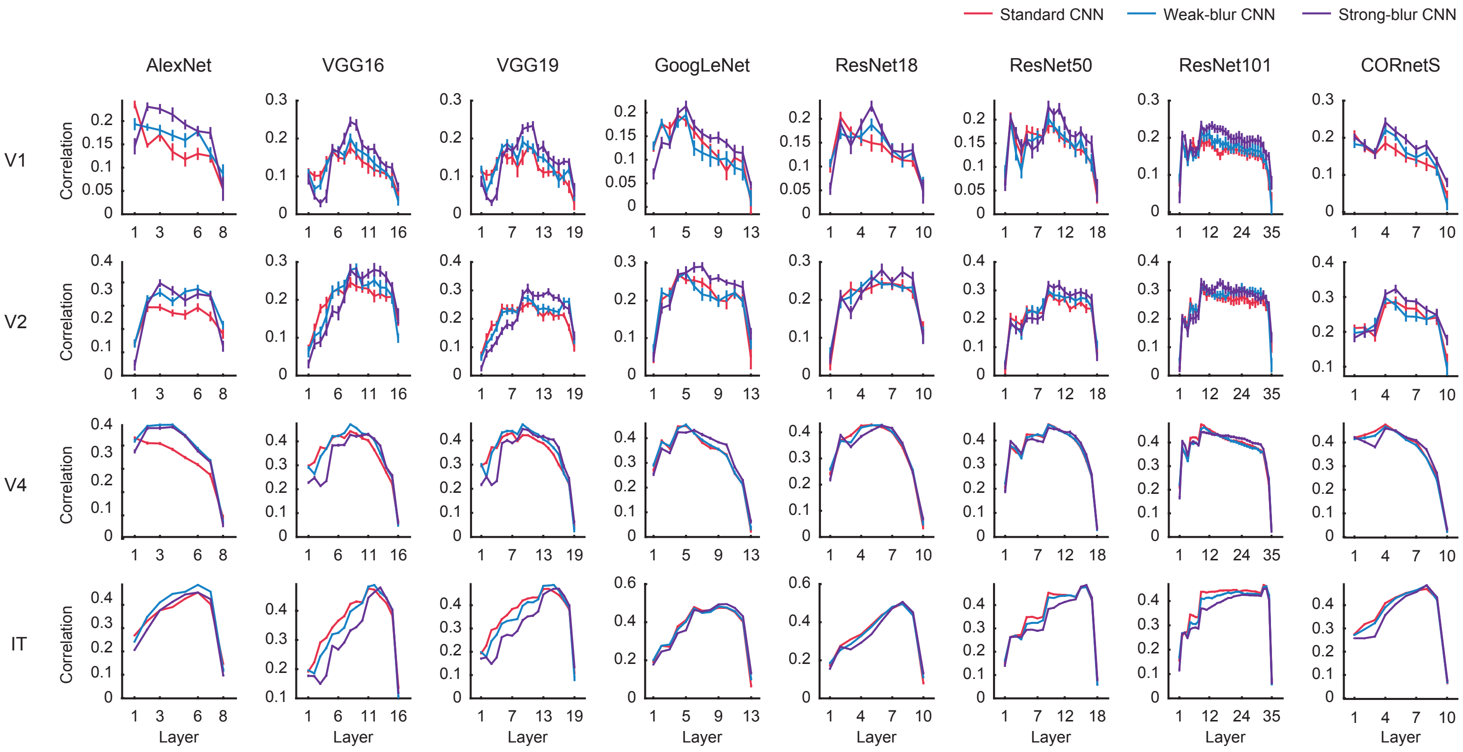


**Supplementary Figure 2.** Correlation between predicted and actual neuronal responses recorded from monkey V1, V2, V4 and IT, plotted for every CNN model by layer. Standard (red), weak-blur (blue) and strong-blur (purple) CNN analysis results are shown on each plot. Data from Schrimpf et al. (2020).

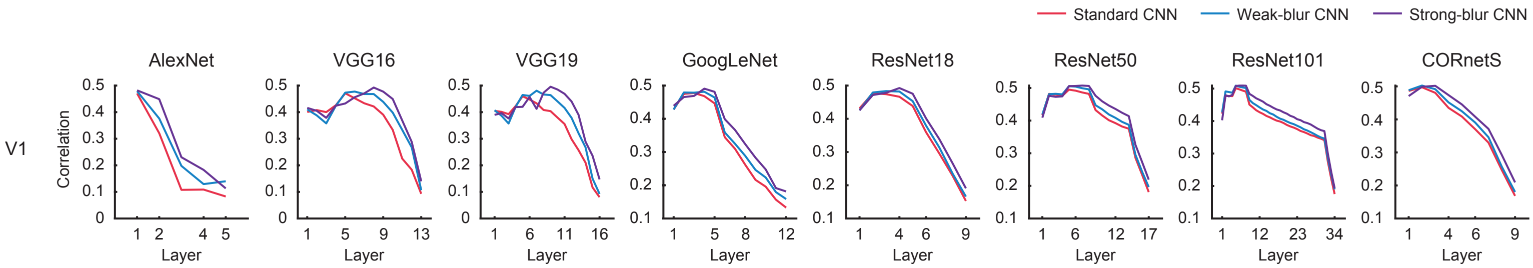


**Supplementary Figure 3.** Correlation between predicted and actual neuronal responses in macaque V1 to thousands of natural and synthetic images. Data obtained from Cadena et al. (2019).

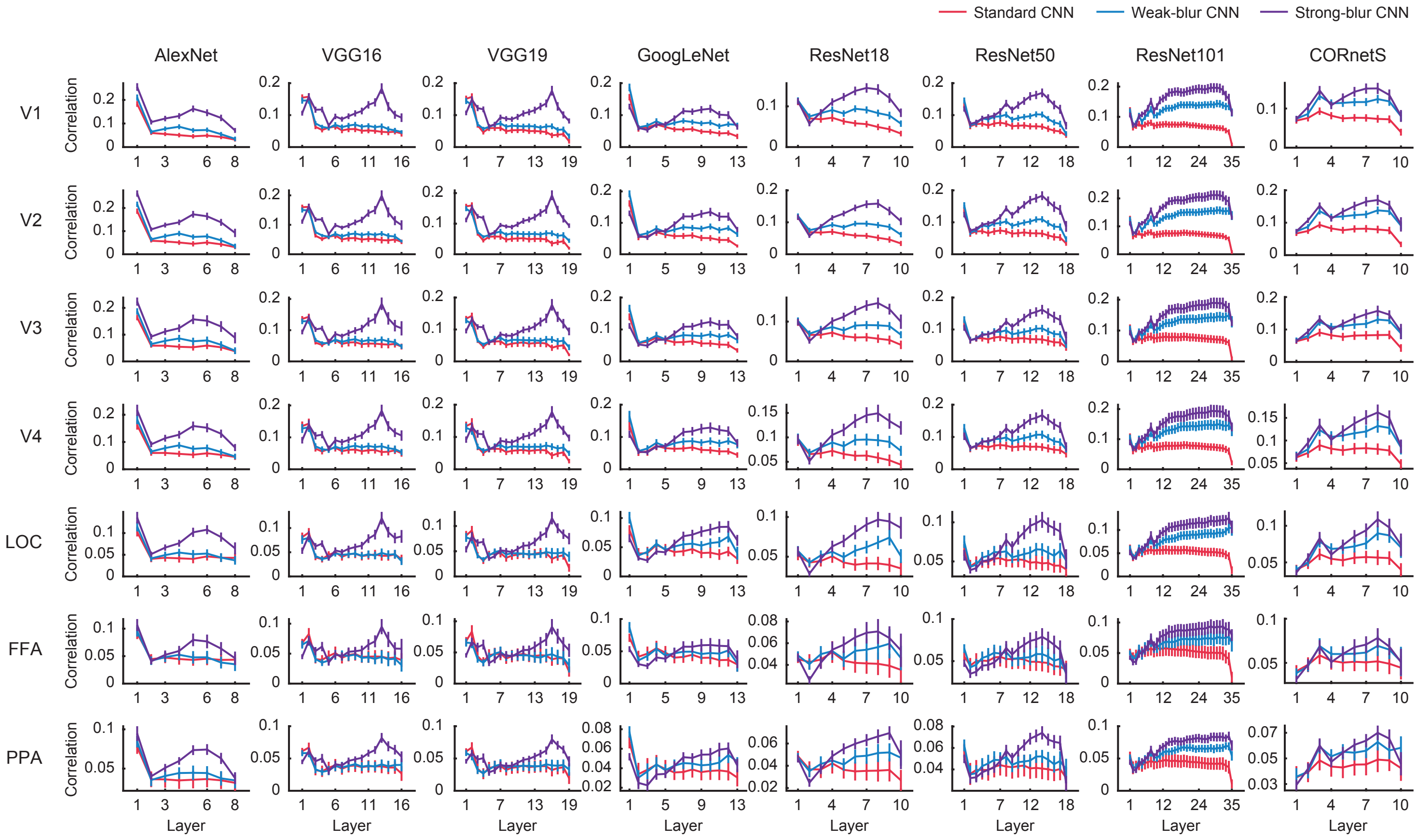


**Supplementary Figure 4.** Layerwise correlation of the RSA matrices between human observers and CNNs across brain regions in all viewing conditions combined (Jang et al., 2021). Standard CNNs (red), weak-blur CNNs (blue), and strong-blur CNNs (purple) were analyzed.

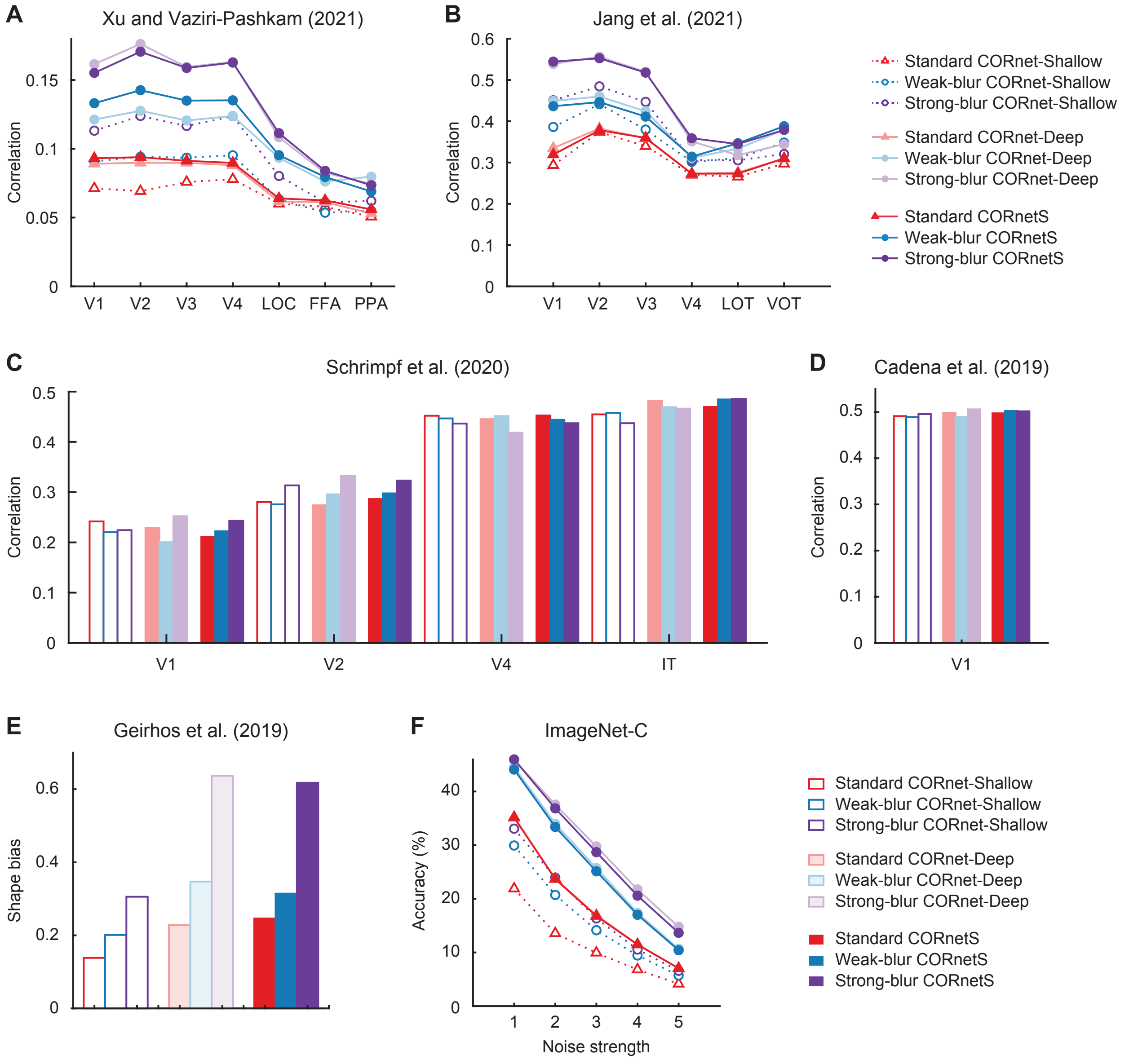


**Supplementary Figure 5.** Evaluation of blur training with a comparative analysis of recurrent neural network CORnet-S, control network CORnet-Shallow that lacks recurrent processing, and another feedforward control network, CORnet-Deep, which performs a matching number of non-linear operations as CORnet-S. Data obtained from **A** Xu and Vaziri-Pashkam (2021), **B** Jang et al. (2021), **C** Schrimpf et al. (2020), **D** Cadena et al. (2019), **E** Geirhos et al. (2019), and **F** ImageNet-C, Hendrycks and Dietterich (2019).

**Supplementary Table 1.**

| CNN architecture | Number of sampled layers | Names of sampled layers |
|---|---|---|
| AlexNet | 8 | features_1, features_4, features_7, features_9, features_11, classifier_2, classifier_5, classifier_7 |
| VGG16 | 16 | Features_1, features_3, features_6, features_8, features_11, features_13, features_15, features_18, features_20, features_22, features_25, features_27, features_29, classifier_1, classifier_4, classifier_7 |
| VGG19 | 19 | features_1, features_3, features_6, features_8, features_11, features_13, features_15, features_17, features_20, features_22, features_24, features_26, features_29, features_31, features_33, features_35, classifier_1, classifier_4, classifier_7 |
| GoogLeNet | 13 | conv1, conv2, conv3, inception3a, inception3b, inception4a, inception4b, inception4d, inception4e, inception5a, inception5b, fc_1 |
| ResNet18 | 10 | relu1, layer1_0, layer1_1, layer2_0, layer2_1, layer3_0, layer3_1, layer4_0, layer4_1, fc_1 |
| ResNet50 | 18 | relu1, layer1_0, layer1_1, layer1_2, layer2_0, layer2_1, layer2_2, layer2_3, layer3_0, layer3_1, layer3_2, layer3_3, layer3_4, layer3_5, layer4_0, layer4_1, layer4_2, fc_1 |
| ResNet101 | 35 | relu1, layer1_0, layer1_1, layer1_2, layer2_0, layer2_1, layer2_2, layer2_3, layer3_0, layer3_1, layer3_2, layer3_3, layer3_4, layer3_5, layer3_6, layer3_7, layer3_8, layer3_9, layer3_10, layer3_11, layer3_12, layer3_13, layer3_14, layer3_15, layer3_16, layer3_17, layer3_18, layer3_19, layer3_20, layer3_21, layer3_22, layer3_22, layer4_0, layer4_1, layer4_2, fc_1 |
| CORnet-S | 10 | V1.output, V2.output.0, V2.output.1, V4.output.0, V4.output.1, V4.output.2, V4.output.3, IT.output.0, IT.output.1, decoder.output |
